# Middle-schoolers’ reading and processing depth in response to digital and print media: An N400 study

**DOI:** 10.1101/2023.08.30.553693

**Authors:** Karen Froud, Lisa Levinson, Chaille Maddox, Paul Smith

## Abstract

We report the first use of ERP measures to identify text engagement differences when reading digitally or in print. Depth of semantic encoding is key for reading comprehension, and we predicted that deeper reading of expository texts would facilitate stronger associations with subsequently-presented related words, resulting in enhanced N400 responses to unrelated probe words and a graded attenuation of the N400 to related and moderately related words. In contrast, shallow reading would produce weaker associations between probe words and text passages, resulting in enhanced N400 responses to both moderately related and unrelated words, and an attenuated response to related words. Behavioral research has shown deeper semantic encoding of text from paper than from a screen. Hence, we predicted that the N400 would index deeper reading of text passages that were presented in print, and shallower reading of texts presented digitally.

Middle-school students (*n* = 59) read passages in digital and print formats and high-density EEG was recorded while participants completed single-word semantic judgment tasks after each passage. Following digital text reading, the N400 response pattern anticipated for shallow reading was observed. Following print reading, the N400 response pattern expected for deeper reading was observed for related and unrelated words, although mean amplitude differences between related and moderately related probe words did not reach significance. These findings provide evidence of differences in brain responses to texts presented in print and digital media, including deeper semantic encoding for print than digital texts.

## Introduction

The use of digital platforms for delivery of instruction and information at school and at home is now requisite for students at all levels, from elementary school through higher education. The increased use of digital materials alongside paper-based materials in learning environments has motivated research into the efficacy of reading and learning in one format versus the other (e.g., [1–5]), and although there is an overall finding for a paper-based advantage, the outcomes have been nuanced. Some reports have indicated no differences between print and digital media with respect to reading ability [1, 6–13], or reading rates and eye movements as measured by eye-tracking [14]. Some authors reported faster reading times in digital compared to paper environments [15, 16], while others reported the reverse [17–20]. Notably, those reporting shorter reading times for computer-based reading also reported a decrease in reading comprehension accuracy in this medium. However, Kim and Kim [20] found that teenagers read faster in the paper-based condition compared to a digital format with a scrolling feature, and also that they scored significantly higher on exams when they studied via paper-based texts. Others [1, 8, 21] reported no difference in reading times between the two media but observed higher comprehension scores in the paper-based condition, suggesting a metacognitive moderating factor. This proposition is supported by results showing that outcomes are poorer on computer-based exams when time is constrained, in contrast to self-paced exams, perhaps because students find it more difficult to self-regulate, monitor task progress, and manage goals and time in digital space [4, 22, 23]. Lauterman and Ackerman [24] also found that the media preferences of exam-takers correlated with performance, suggesting that findings for a computer-based inferiority may be associated with a deficit in the knowledge and skills necessary to navigate in the digital medium.

Reading comprehension seems also to be moderated by both depth of remembering and text genre. Comprehension scores measured by understanding the gist of what was read have been repeatedly shown not to differ between narrative and expository texts, regardless of the medium of text presentation [1, 2, 6, 12]. In contrast, reading both expository and complex texts from paper seems to be consistently associated with deeper comprehension and learning [1, 2, 25]. Mangen et al. [26] observed an advantage for print over digital media for both narrative and expository texts.

These varied outcomes may be attributed to a number of factors, such as differences in age and grade-level of study participants, their learning goals, and learned strategies. For elementary students, medium of presentation has been shown to have little influence on comprehension of simple texts [6, 7]. Lenhard et al. [15] found that elementary and middle school children were faster at completing a reading comprehension assessment on computer compared to paper under time constraints, but at the expense of accuracy. Critical reading skills of high school and college students were compared by Eshet-Alkalai and Geri [27], who found that younger students performed better when reading news in digital formats compared to paper, while college students performed better on the same task when reading in paper formats.

Against this lack of clarity in the behavioral findings, there has been little brain imaging work to further elucidate the mechanisms that underpin reading in print versus digital formats. Kretzschmar et al. [14] recorded electroencephalography (EEG) during their eye-tracking paradigm, that was designed to evaluate whether stated preferences for the printed medium (versus one of two digital devices) correlated with indices of text engagement in young and older adults. Comprehension accuracy did not differ with text presentation medium for either group, but the older adults showed shorter mean fixation durations and lower EEG theta band voltage density when reading from a tablet computer in comparison to an e-reader or a printed page. Younger adults did not show any such differences, and Kretzschmar et al. interpret the observed differences as relating to limitations on memory encoding and retrieval for the older adults, affected by reduced contrast sensitivity, that could be somewhat ameliorated by the backlit display of the tablet computer. However, there exist currently no other reports of EEG measures applied to the question of reading in different media, and crucially there have been no investigations of brain responses to print vs. digital text processing in children.

For our investigation, we drew upon depth of processing theory, first posited by Craik and Lockhart [28]. The premise of this theoretical framework is that shallow information processing yields less durable episodic memory traces, while deeper processing results in more durable traces. The central claim is that the more deeply information is processed, the more durable the associated memory traces. Kintsch [29, 30] has described text comprehension as a dynamic process of constructing meaning from semantic relations among words in the text and stored knowledge about subject matter. According to seminal work by Craik and Tulving [31], processing of verbal text information requires the use of semantic processes (protocols concerning the ways in which words work together to create meaning); hence, text processing strategies for reading may involve drawing on contextual, semantic, grammatical, and phonemic knowledge in systematic ways to work out what information is conveyed by a text. Such strategies would allow an encoded unit to be integrated with knowledge of the world or “semantic memory” (e.g., [32]). At retrieval, informational cues would then tap into this semantic memory structure to reconstruct an initial encoding [31].

Based on this theoretical framework, we proposed that *the medium* whereby readers engage with text/reading material would be a crucial determinant of differences in depth of processing, and consequently the durability of the semantic memory structure that is established. Congruous encoding between a semantic structure already established by a reader and a semantic structure associated with a newly encoded unit should facilitate efficient comprehension of a text, first because a meaning-referenced elaborated trace network is formed, and second because robust congruent semantic encoding also entails alignment with the structure, rules, and organization of semantic memory [31, 33].

Consistent with this view of semantic structure and encoding processes, we hypothesized that depth of semantic encoding is key for reading comprehension and for congruency between existing semantic structures and the semantic structures encoded by probe words. Based on previous research, semantic encoding of text presented on paper is deeper than that of text presented digitally [1, 26]. Therefore, our experimental approach to measuring reading comprehension in the brain made use of a signature of electrophysiological activation associated with semantics in language processing: the N400 event-related potential (e.g., [34, 35]).

The N400 event-related potential (ERP) indexes brain response differences between expected and unexpected stimuli. Since we hypothesized that the encoding of word meaning during the reading experience is critical for comprehension, then we should be able to index shallow vs. deep information processing of text delivered in print or digitally by observing differences in N400 responses to probe words that were selected to be related, moderately related, or unrelated in meaning to written passages. Based on this hypothesis and given that both the culturally prevailing view and data meta-analytic studies [1–5] suggest that reading digitally presented text promotes shallower engagement than print, our predictions for the electrophysiological index were as follows: 1) In the digital reading condition, the N400 amplitude response to related word probes was predicted to be attenuated compared to moderately related and unrelated word probes, with amplitude differences between moderately related and unrelated word conditions expected to be equivalent; and 2) In the print reading condition, the N400 amplitude response to the three conditions is predicted to be graduated. Specifically, the amplitude measures were predicted to increase in their negativity such that the response to the related words would be most attenuated, followed by the moderately related words, with words that are unrelated to the text passage eliciting the greatest negativity. Differences in the N400 ERP response between the two mediums for the moderately related word conditions may offer essential insights about the neurocognitive processing underlying reading comprehension, and whether readers in some situations process text somewhat more shallowly under conditions of digital text presentation than when processing text via print presentation.

## Materials and methods

### Participants

We collected data from 65 participants from the New York City metropolitan area and were able to retain data from 59 (five were removed due to unusable behavioral data; one was removed due to low numbers of EEG trials per condition following artifact detection – detailed further below).

The mean age of retained participants was 10.88 years (*SD* = 0.77); of these, 28 identified as male and 28 as female, with one participant giving no response to this question. Most participants were in 5th (*n* = 21) or 6th grade (*n* = 22) at the time of their lab session, as expected; the remainder were in 4^th^ (n = 2), 7^th^ (*n* = 10), or 8th grade (*n* = 2), and two indicated “other”. All participants were from households with at least one parent or guardian who attended some post-secondary education, with the majority having earned degrees: associate degree (3.5%), bachelor’s or undergraduate degree (28.1%), master’s degree (52.6%), or doctorate (10.5%). Household annual income was reported as $150,000 per year or above for 56% of participants, with the balance of participants spread among the other income brackets (no response; $35,000 – $49,999; $50,000 – $74,000; $75,000 – $99,999; $100,000 – $149,999).

### Stimuli

#### Passages

Based on the key finding that a paper-based reading advantage is seen largely in studies using informational or a mix of informational and narrative text [1, 2, 25], all reading passages were developed as informational texts. Several additional goals were set for the passage development so that passages could be used as controlled experimental stimuli yet remain similar to text that might be found in a classroom setting. The passages covered a range of topics to account for differing interests among participants. We also controlled the level of reading difficulty and complexity while maintaining grade-level and age-appropriate standards. Finally, we ensured that there was sufficient content for generation of word probe stimuli for the subsequent single-word semantic relatedness judgement task. These passages were limited to relatively simple sentence structures (minimizing relative or subordinate clauses) while preserving the historical and scientific accuracy of the presented material.

Eight passages were created in thematic pairs to allow for later comparison across mediums. The passages were matched for length with respect to average number of words per sentence (*mean* = 11.736, *SD* = 1.073), number of total words (*mean* = 189.125, *SD* = 9.250), and number of sentences (*mean* = 16.250, *SD* = 1.389). Readability scores were calculated and matched for each passage, specifically the Flesch-Kincaid Grade Level ([36]: *mean* = 5.775, *SD* = 0.711), Gunning Fog score ([37]: *mean* = 7.950, *SD* = 0.795), and the SMOG index ([38]: *mean* = 6.388, *SD* = 0.541). In addition, we matched the passages on Propositional Count (PC), a quantification of the number of semantic units and their connections within the text ([39–41]: *mean* = 65.750, *SD* = 2.188).

#### Passage Reading Comprehension Measure

To assess participant comprehension for each text, it was necessary to develop passage-specific assessments. The Sentence Verification Technique (SVT; [42]) is an assessment procedure based on the theoretical assumption that reading comprehension is a constructive process involving interactions between incoming discourse and the reader’s prior knowledge structure. SVT comprehension test items are graded questions derived from texts that require varying levels of passage knowledge to answer. The four question types specified within the framework are: *Explicit/Original*, whereby a sentence directly from the text must be identified as such by the reader; *Paraphrase,* whereby a sentence from the text is paraphrased, and must be identified as such; *Meaning Change,* a sentence that changes an aspect of meaning presented in the text, and which should therefore be rejected by the reader; and *Unrelated/Distractor* items. We used *Explicit* and *Unrelated* categories from the SVT framework as defined but made adaptations to the other two question types. For the *Meaning Change* condition, we altered sentence meanings by replacing only a single propositional predicate with a related probe word. Our *Paraphrase* items were not sentences from the passage themselves, but true statements that combined propositions from across the entire text. SVT sentences were constructed to minimize syntactic complexity (active sentences only, no subordinate clauses), matched for sentence length (mean 10.625 words per sentence, *SD* = 1.619), and controlled with respect to the age of acquisition (AoA) of individual words (based on ratings from [43]; mean AoA for all SVT items = 5.984, *SD* = 1.936).

Conventionally, SVT items elicit a binary response (*Yes, No*) making scoring a simple process. For our purposes, we provided students with three selection options based on the relatedness of the sentence to the passage: (1) *I read exactly this sentence in the passage*; (2) *The facts in this sentence were in the passage*; or (3) *None of the facts in this sentence were in the passage*. We applied a binary scoring procedure to *Explicit* and *Distractor* responses to SVT items: an *Explicit* item was scored correct if response (1) was selected, and a *Distractor* item was scored correct if response (3) was selected. In the conventional SVT framework, *Meaning Change* test items should all be identified as false, whereas our items were a mixture of true and false statements. Per convention all *Paraphrase* items were true. For analyses, we marked a *Paraphrase* item as correct if a respondent indicated response (2); we scored the *Meaning Change* items as correct if either (2) or (3) was chosen, depending on the assigned truth value for that statement.

#### Validation of Passages and Passage Comprehension Items

Prior to conducting the experiment, we collected online reader response data to the eight passages via Panelbase LLC (panelbase.net). These data ensured that stimuli were balanced with respect to the following parameters: reading time for each passage; participant interest in the passages; self-reports of reading difficulty; a set of cloze questions to evaluate attention to each passage; and the constructed SVT items. Respondents represented a random sample of students matching the study target population, drawn from U.S. urban areas excluding New York City. Between 70 and 80 participants completed a survey that included a selection of two of the eight passages. The results of this pre-study validation procedure pointed to general equivalency across these eight passages in terms of difficulty and accessibility, as well as general responses to the SVT question types. Analyses of this data identified that one passage set (two thematically related passages) was more difficult relative to the others, and so these two passages were excluded from the experiment.

#### Stimulus Probe Words

Probe words for the semantic relatedness judgment paradigm were generated by identifying verbs or nouns at the center of propositions in each passage as targets for semantic field interrogation. Using the WordNet 3.0 database [44–46], each selected verb and noun was used as a search term and the relevant propositional sense was identified in the returned synset listings. Each synset was then expanded and lexical items (uninflected, nonderived) of the same word class as the target proposition were selected from synset lists. Frequency (Zipf scores: [47]), age of acquisition (AOA: [43]) and length characteristics (NLET, NPHON, and NSYLL, all from the MRC Psycholinguistics Database: [48]) were determined for each item. Items were included in the semantic rating experiment only when their frequency and AOA ratings were within 1 SD of the mean for the target passage. Concreteness was also evaluated, and there were no differences between any of the word conditions with respect to that property [49]. Probe words fell into three categories: *Related*, *Moderately Related*, and *Unrelated*.

Potential items for the *Related* category of word probes were identified based on the specific propositions identified in each passage. Based on the number of related words that were identified per passage, a number of words from a pool of semantically unrelated words that were likewise matched on AOA and frequency were also included, to yield up to 100 target items per passage. Relatedness ratings were validated using the online platform Prolific (prolific.co). Semantic ratings were solicited from an adult population as opposed to the target population of middle-school students given that adults are more likely to have well-developed semantic networks [50]. Adult raters read each text passage and then rated candidate probe words for relatedness to the passage on a scale from 0% to 100%. Each participant rated potential probe words for two passages. For each passage, the candidate probe words were rated on relatedness by 100-150 participants. Ratings were trimmed to remove ratings of 0% or 100% and Gaussian mixture modeling (e.g., [51, 52]) was applied to data for each passage to cluster ratings into the three stimulus categories: related, moderately related and unrelated. The 20 words closest to the mean score within a cluster were assigned to that category; if a category contained fewer than 20 words, only that many words were assigned. Probe words that applied to multiple passages were assigned to the category and passage for which they were closest to their cluster mean, and the next closest word was chosen for the other passage.

The moderately-related words, those that fell within the center cluster, were labeled as “chimera” items, reflecting the possibility that a word that is moderately related to some context could also be identified as moderately unrelated to that context. The judgement task for N400 elicitation required a binary decision concerning relatedness (related vs. unrelated), and these items were evaluated as somewhere in between. The chimera words were crucial to our predictions, as we anticipated that deeper processing would facilitate participants’ identification of chimera words as related to the preceding textual context, while shallower processing would be more likely to result in identification of chimera words as unrelated.

#### Psychometric Measures

All participants completed a set of standardized assessments, plus an additional assessment of auditory working memory, as follows:

- Wechsler Intelligence Scale for Children V [53] Digit Span subtest – an assessment of working memory capacity
- Woodcock Reading Mastery III [54] Passage Comprehension subtest – an assessment of general reading comprehension
- Woodcock reading Mastery III [54] Word Attack subtest – an assessment of phonemic decoding ability
- Swanson Listening Sentence Span Task (LSST; [55]) – an assessment of working memory span that is mediated by language

### Data Collection

Data were collected in three phases: Phase 1 for the online administration of psychometric assessments; Phase 2 for the EEG recordings and the immediate passage recall comprehension measure; and Phase 3 for the online administration of the passage retention comprehension measure. All informed consent and experimental procedures were carried out with approval of the Teachers College, Columbia University Institutional Review Board (Protocol # 22-173). Written informed consent/assent was obtained from all individual participants included in the study.

#### Phase 1: Psychometric Assessments

In Phase 1, responses to behavioral assessments were collected by two trained assessment administrators online during video conference sessions. The parent/guardian received a study overview, consent, and assent forms in advance of the scheduled appointment. They were asked to select a quiet setting with the home with minimal distractions where the participant could complete the assessments. Online, the assessment administrator reviewed the materials and responded to questions before the parent/guardian and their child completed the consent and assent forms obtained via a Qualtrics survey. The sessions were approximately 25 to 30 minutes in length and audio recordings were stored for the purpose of second-scoring of measures.

#### Phase 2: EEG Recording

In Phase 2, participants and their accompanying parent/guardian attended the Neurocognition of Language Lab at Teachers College, Columbia University. High-density EEG data were continuously recorded in NetStation 4.3.1, using a 128-channel HydroCel Geodesic Sensor Net (MagStim Electrical Geodesics, Inc.). Signals were amplified using a NetAmps 200 series amplifier. Samples were collected at a rate of 500 Hz; an online low-pass filter of 200 Hz and high-pass filter of .1 Hz were applied. Impedances were kept below 40 kiloohms and were re-checked between blocks. Participants completed sessions in an electrically shielded and sound-attenuated room, seated 65 cm from a computer monitor with a brightness of 75 cd/*m^2^*.

Each participant was first exposed to texts that were presented via either a paper booklet (print) or a laptop screen (digital). For the digital reading condition, visual readability variables (contrast, brightness, text size) between the laptop screen and the stimulus presentation screen were held equivalent. The order of medium and passage presentation was balanced between participants. Passage reading time was recorded, and then participants completed two tasks, presented using E-Prime 3.0 (Psychology Software Tools, LLC).

First, participants read one text passage in their assigned medium. Then they completed the semantic relatedness judgment task in response to single probe words presented on a computer screen. They responded to each word by pressing one button to indicate that a word was related to the passage, and another if they thought the word was unrelated (see Fig 1).

After the semantic relatedness judgment task, the SVT recall comprehension test items were displayed, and participants were prompted to respond. This procedure was repeated for two additional passages in the selected medium (either print or digital). Then, the medium of presentation was switched, and the process was repeated for another three passages.

**Fig 1.**
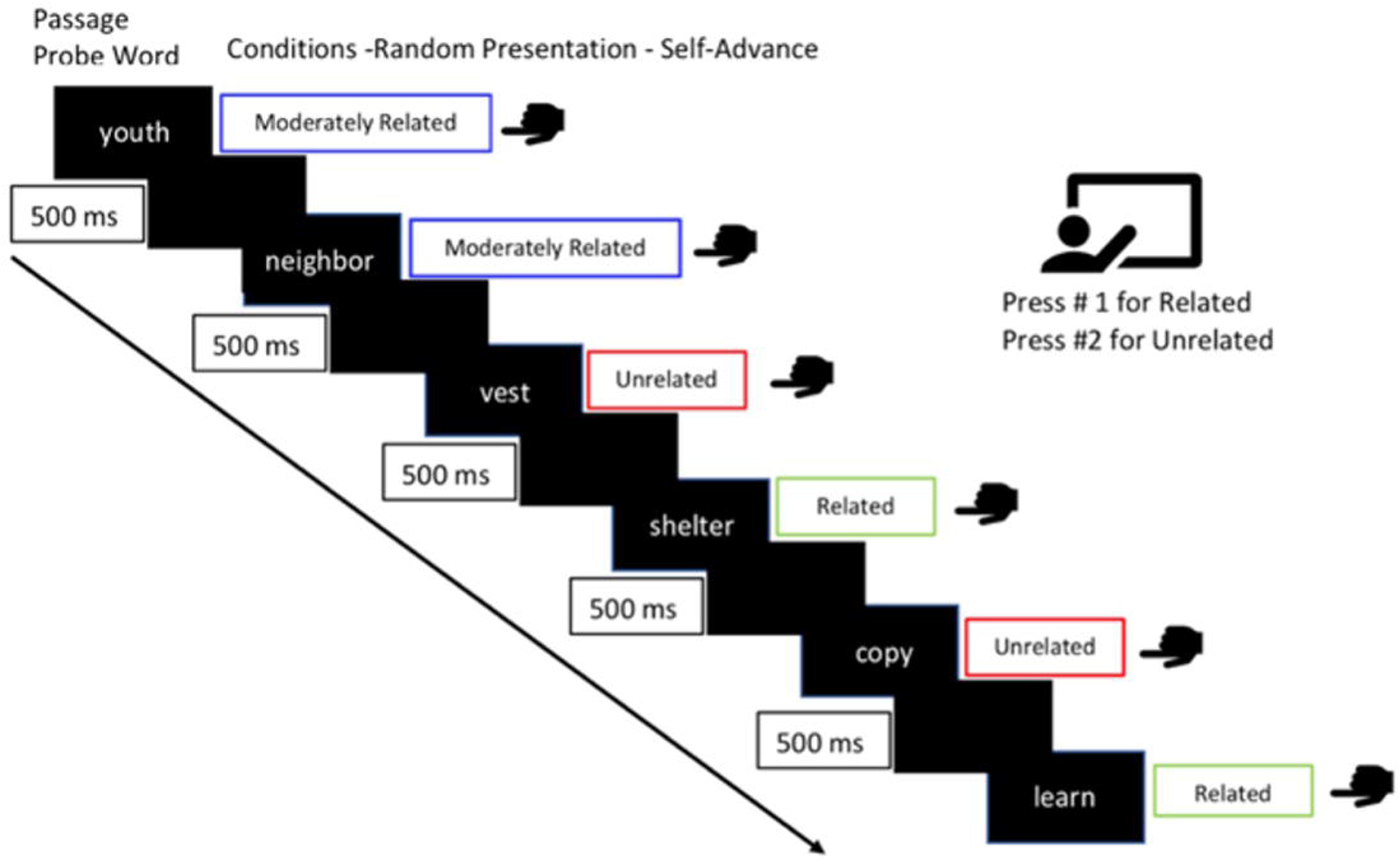
Timeline for example trials from the semantic relatedness task.

#### Phase 3: Passage Retention Measure

Following the EEG recording session, participants were emailed a link to the follow-up Qualtrics retention comprehension survey. This consisted of the same SVT recall comprehension test items that they completed during the EEG portion of the study and was included to provide an indication of information retention. Participants were asked to complete the measure within 24 hours of their visit to the lab, but survey responses were accepted up to seven days after their lab visit.

### EEG Data Analysis

#### Pre-Processing

EEG data were pre-processed using the Harvard Automated Processing Pipeline for Electroencephalography (HAPPE; [56]), specifically the event-related extension (HAPPE+ER; [57]). The sensitivity of the HAPPE procedures allows for more trials to be kept and averaged when dealing with high-variance data such as those associated with children. Globally bad channels were detected and removed from the remainder of the pipeline. Across all participants, an average of 93.6% (SD: 4.4%) of channels were good, with a range of 61.2% to 99.2%. A hard wavelet threshold was applied to remove artifacts from the continuous EEG data, a technique that improves upon previous methods of detecting artifacts to retain more of the EEG signal instead of rejecting segments at this stage [57]. A pre-established bandpass filter from 0.1-40 Hz was utilized, and data were segmented from 100 milliseconds (ms) before stimulus presentation to 750 ms post-presentation.

Segmented data were subjected to baseline correction, whereby the average of the EEG recorded during the baseline period for each epoch was subtracted from the post-stimulus period. Bad data within each segment were interpolated and segments were rejected based on a joint probability criterion as well as amplitude cutoffs of -150 and 150 microvolts. Globally bad channels were replaced based on spherical spline interpolation of data from surrounding electrodes, and data were re-referenced offline to the average of the left and right mastoid channels (electrodes 57 and 100).

Participants were excluded from further analysis if more than 40% of trials for any passage were rejected. Of 65 participants, one was excluded due to low numbers of trials in the final analysis and others due to inability to use behavioral data; analyses were therefore based on data from 59 participants. For all retained participants, at least 50% of trials were deemed usable; on average, 66.5% of trials were usable (SD: 5.8%; range: 53.2% to 82.3%). The numbers of trials per participant did not vary significantly across medium or passage. For the related and unrelated conditions, error trials (trials in which a participant had misidentified a related word as unrelated, or vice versa) were also excluded from further analysis. All trials were kept for the chimera condition, as their intermediate level of relatedness makes them hard to accurately categorize in a binary fashion. In the print medium, 910 related trials, 2,024 chimera trials, and 1,961 unrelated trials were used in subsequent analyses; in the digital medium, trial numbers came to 909 related trials, 2,016 chimera trials, and 2,028 unrelated trials.

Baseline-corrected epochs for each word condition were then averaged together for each individual participant, providing individual averages per medium and condition. Individual event-related potentials were interrogated for mean amplitude of the target component within an *a priori-*established time window, 300-500 milliseconds post-stimulus. Individual averages per condition were then grand averaged to generate group ERP waveforms.

#### Montaging

N400 montages vary across studies (e.g., [58]). The electrode montage for investigation of the N400 component was selected based in part on the N400 context and discourse literature [59–65]. Fig 2 below indicates the montage of interest; all plots of the derived event-related potentials relate to this montage.

**Fig 2.**
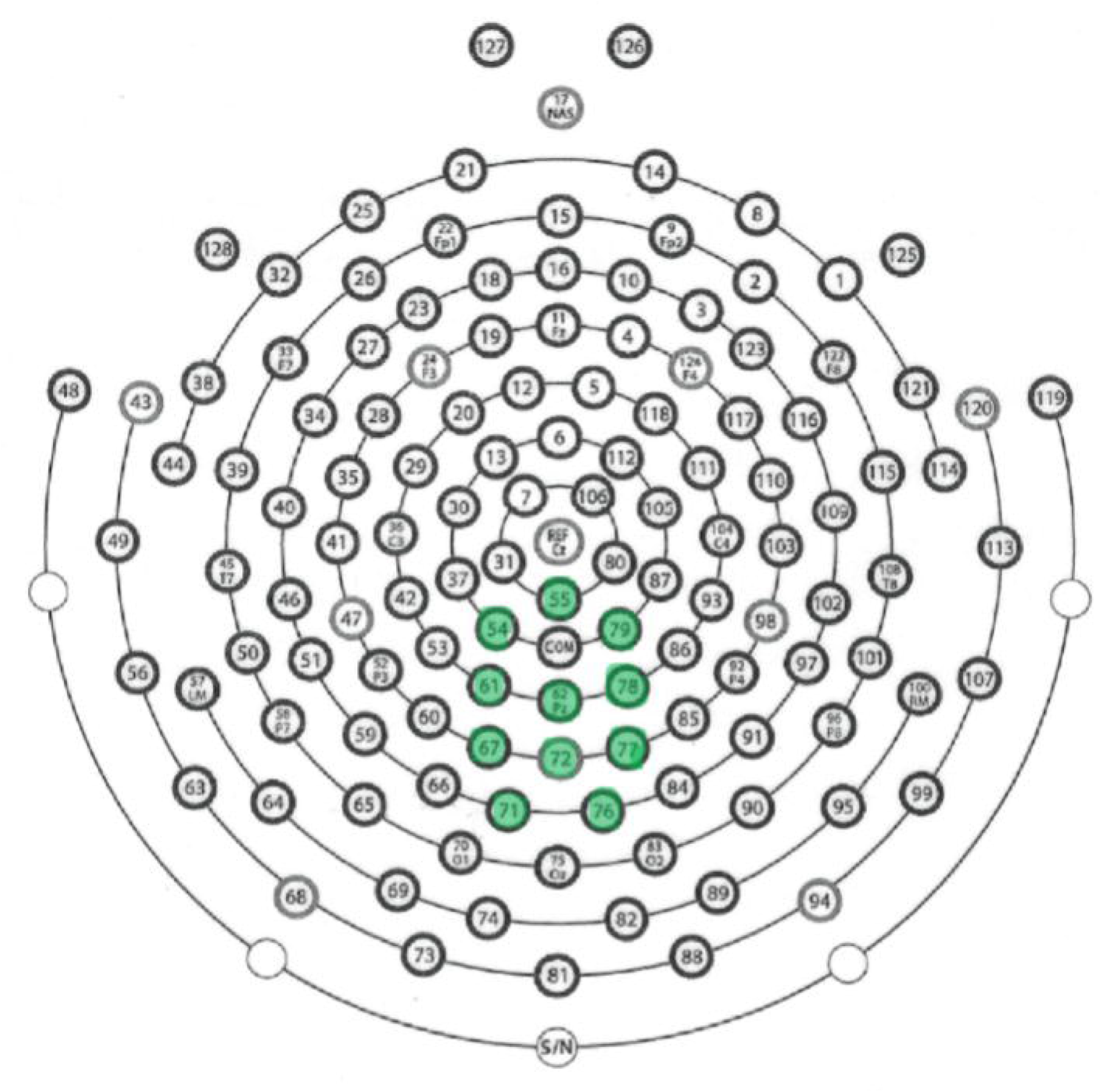
Montage for N400 analysis. Electrodes included in the analysis montage are indicated in green: electrode numbers 54, 55, 61, 62, 67, 71, 72, 76, 77, 78, 79.

## Results

### Phase 1: Psychometric Assessments

All participants completed a set of standardized assessments, plus an assessment of auditory working memory. Table 1 below provides mean scores and standard deviations for each assessment for all included participants (*n* = 59). We applied a criterion to include only those participants whose scores on all assessments were within 3 standard deviations of the sample mean for each assessment. All participants met this criterion. While we did not have any outliers that needed to be removed from the data analysis, a range of abilities was represented within this sample of middle school students.

**Table 1.** Mean scores and standard deviations for psychometric reading assessments.

| Source Assessment Battery | Subtest / Scoring Sample | Mean standard score (SD) |
| --- | --- | --- |
| WISC-V | Digit Span – forwards | 11.49 (3.28) |
|  | Digit Span – backwards | 10.93 (3.37) |
| WRMT-III | Word Attack – by grade | 107.20 (11.89) |
|  | Word Attack – by age | 107.90 (11.70) |
|  | Passage Comprehension – by grade | 116.05 (14.44) |
|  | Passage Comprehension – by age | 117.81 (14.76) |
| Listening Sentence Span Task | N/A | 2.10 (1.34) |

We evaluated correlations between these measures to determine which assessments of working memory (digit span and LSST) were correlated with the measures of language skill from the WRMT-III. A review of the relationships between different measures revealed no significant correlations between the Listening Sentence Span Task (LSST) and traditional working memory measures (forward and backward digit span: *r* = .088, *p* =.507 and *r* = .155, *p* = .241, respectively). However, there was a significant correlation between the digit span scores (*r* = .559, *p* < .001). The LSST measure was found to be positively correlated with both word attack and passage comprehension scores (see table 2, below).

**Table 2.**
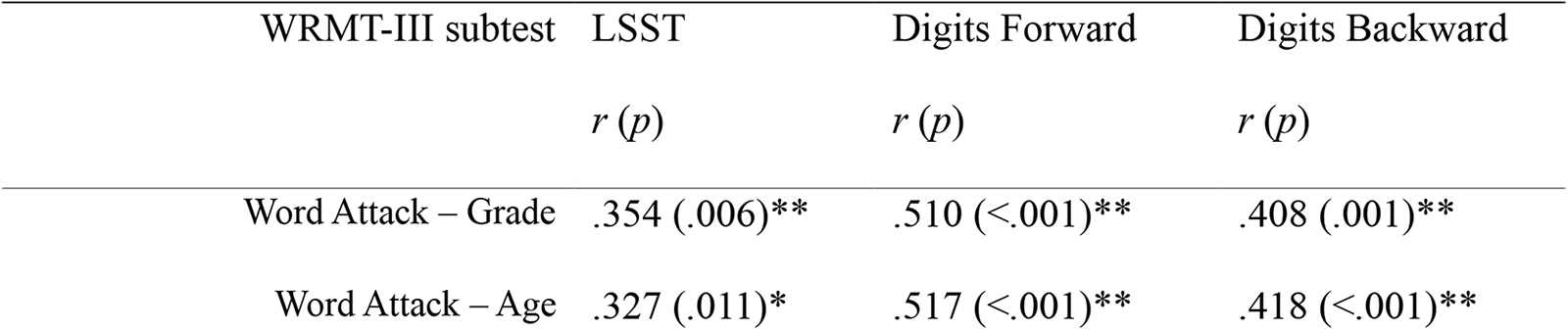
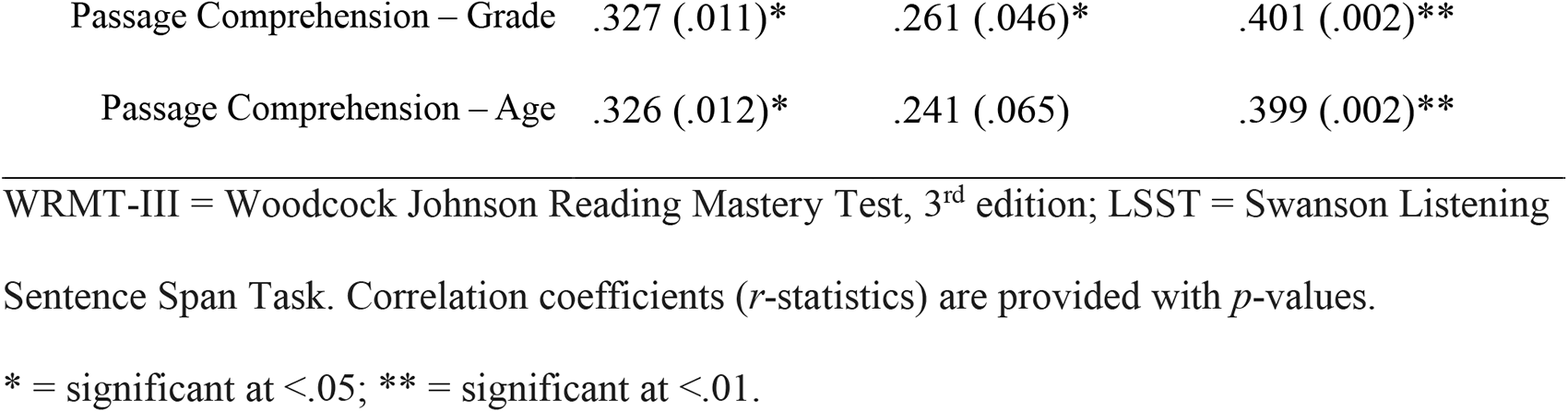
Correlations between scores on working memory and language assessments.

| WRMT-III subtest | LSST | Digits Forward | Digits Backward |
| --- | --- | --- | --- |
| | $r(p)$ | $r(p)$ | $r(p)$ |
| Word Attack – Grade | .354 (.006)** | .510 (<.001)** | .408 (.001)** |
| Word Attack – Age | .327 (.011)* | .517 (<.001)** | .418 (<.001)** |
| Passage Comprehension – Grade | .327 (.011)* | .261 (.046)* | .401 (.002)** |
| Passage Comprehension – Age | .326 (.012)* | .241 (.065) | .399 (.002)** |

WRMT-III = Woodcock Johnson Reading Mastery Test, 3<sup>rd</sup> edition; LSST = Swanson Listening
Sentence Span Task. Correlation coefficients (*r*-statistics) are provided with *p*-values.
\* = significant at <.05; \*\* = significant at <.01.

These findings indicate that (a) working memory was within a typical range across the group of participants; (b) that working memory is important to control in experimental approaches to reading comprehension; and (c) that working memory is *not* likely to be a factor influencing neurophysiological response differences between passages in this experiment.

### Phase 2: Event-Related Potentials

We examined the grand-averaged N400 responses to all probe word conditions (i.e., related, chimera, unrelated) within each medium (i.e., following texts presented in digital vs. print media). Plots displaying the grand-averaged waveforms of participant responses for each probe word condition within each presentation medium condition are shown in Fig 3. Within each medium condition, paired-samples *t*-tests were conducted to observe differences between the three probe word types. Following text presented in the digital medium, the N400 response to related words was significantly different from the response to both chimera words (mean difference = 1.644 µV, *t* (58) = 3.562, *p* = .0012, *d* = .464) and unrelated words (mean difference = 2.204 µV, *t* (58) = 4.055, *p* < .003, *d* = .528). The response to chimera words was not significantly different than the response to unrelated words in the digital text condition (mean difference = .560 µV, *t* (58) = 1.718, *p* = .138, *d* = .224). Following texts presented in the print medium, the response difference between related and unrelated words was significant (mean difference = 1.043 µV, *t* (58) = 2.755, *p* = .012, *d* = .359). The difference between related and chimera words was not significant (mean difference = .304 µV, *t* (58) = .798, *p* = .642, *d* = .104), but a significant difference between chimera words and unrelated words was observed (mean difference = .739 µV, *t* (58) = 2.546, *p* = .021, *d* = .331). All tests were controlled for multiple comparisons via Bonferroni correction within medium.

**Fig 3.**
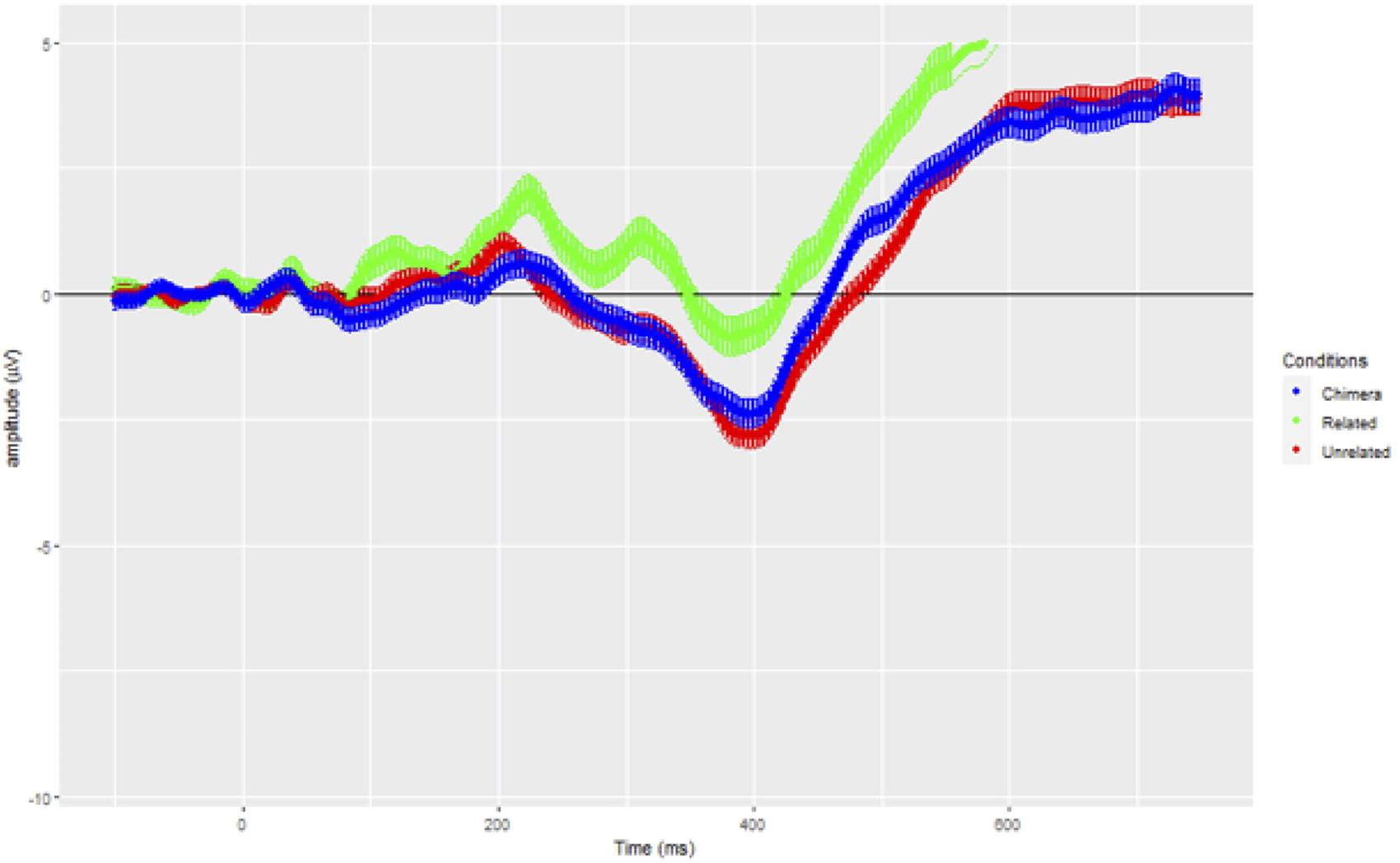
Grand-averaged waveforms in response to the semantic relatedness task following digital text presentation. Includes all retained participants, correct response trials only for related and unrelated word conditions, and all responses to chimera words (no error criterion for this condition). Variance around the mean waveforms is shown as shadow. Green: Related condition; blue: Chimera condition; red: unrelated condition.

**Fig 4.**
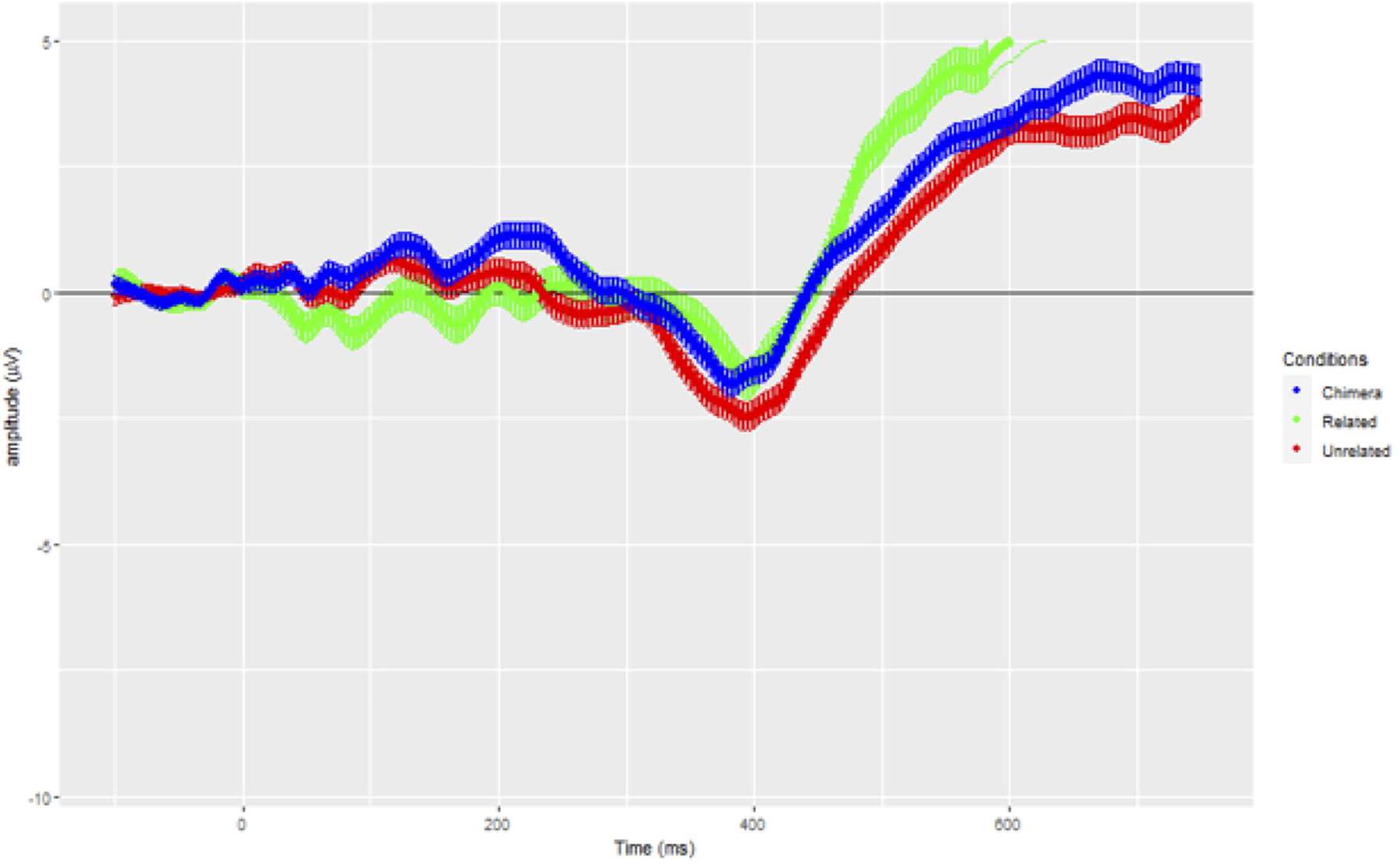
Grand-averaged waveforms in response to the semantic relatedness task following digital text presentation. Includes all retained participants, correct response trials only for related and unrelated word conditions, and all responses to chimera words (no error criterion for this condition). Variance around the mean waveforms is shown as shadow. Green: Related condition; blue: Chimera condition; red: unrelated condition.

### Experimental Results: Behavioral Findings

Following each passage, participants were asked to decide whether each word shown on screen was related or unrelated to the passage they had just read. Related words were both scored as “correctly identified” if the participants indicated they were related to the passage; unrelated words were similarly coded if they were marked as unrelated. For the related words, participants correctly identified on average 45.29% (SD: 21.116) of words as related in the digital medium, and 44.18% (SD: 21.558) in the print medium. For unrelated probes, on average 96.92% (SD: 4.913) of words in the digital medium and 96.68% (SD: 7.561) in the print medium were selected as unrelated to the passage. For chimera words, there was no error criterion; 15.33% (SD: 16.22) of the chimera words were identified as “related” in the digital medium and 15.44% (SD: 14.22) as “related” in the print medium. A two-way repeated measures ANOVA revealed no interaction between medium and category (*F* (1, 58) = .179, *p* = .674) or main effect of medium (*F* (1, 58) = .505, *p* = .480); however, there was a main effect of condition (*F* (1, 58) = 277.261, *p* < .001). Planned comparisons (*t*-tests) confirmed significant differences in accuracy between conditions, with unrelated words being identified significantly more accurately than related words (following text reading in the digital medium: *t* (58) = -16.314, *p* <.001; print medium: *t* (58) = -15.382, *p* < .001).

Reaction times for each word were also recorded for each participant. Following digital text reading, average reaction time for related words was 1,547.064 ms (*SD* = 490.124), for the chimera words was 1,454.827 ms (*SD* = 472.169), and for the unrelated words was 1,352.923 ms (*SD* = 481.730). In the print medium, average reaction time for the related words was 1,502.545 ms (*SD* = 506.423), for the chimera words was 1530.480 ms (*SD* = 559.819), and for the unrelated words was 1,319.922 ms (*SD* = 420.521). A two-way repeated measures ANOVA revealed a significant interaction between medium and category (*F* (2, 116) = 4.278, *p* = .016). There was no significant effect of medium, confirming that reaction times to individual words following reading in print or on a screen did not differ. A significant simple main effect was found for word category (*F* (2, 116) = 23.334, *p* <.001), and planned comparisons (paired-samples t-tests) confirmed that, in the digital medium, reaction times to the related words were significantly longer than to either the chimera (*t* (58) = 3.352, *p* < .001) or the unrelated words (*t* (58) = 4.922, *p* < .001); however, reaction times did not differ significantly between chimera and unrelated words (*t* = 2.684, *p* = .005). In the print medium, reaction times to the related and chimera words were both significantly longer than to the unrelated words (related vs. unrelated: *t* (58) = 4.389, *p* < .001; chimera vs. unrelated: *t* (58) = 5.216, *p* < .001), but the reaction times were not different between related and chimera words (*t* (58) = -0.790, *p* = .433).

### Comprehension Accuracy

#### Immediate Recall Comprehension Task

The reading of each passage was followed by a set of eight sentence verification items to evaluate participants’ comprehension of the preceding passage. The eight items were of four different types, as described above: explicit, paraphrase, meaning change, and unrelated. These four types of questions were designed to probe different aspects of understanding of the text and different levels of difficulty with respect to recall as well as recognition of ideas and concepts from the texts.

Responses to the sentence verification items were not recorded for 9 of the 59 participants due to software malfunction during data collection. Thus, the results below include data for 50 participants. Accuracy for these items is presented below in Table 3, separated by medium.

**Table 3.** Mean percent correct responses for each sentence verification item type, immediate presentation.

| Sentence Verification | Digital Text Presentation | Print Text Presentation |
| --- | --- | --- |
| Item Type | % Correct (SD) | % Correct (SD) |
| Explicit | 64.33% (21.96%) | 65.33% (20.16%) |
| Paraphrase | 52.33% (27.56%) | 54.00% (22.22%) |
| Meaning Change | 30.00% (14.68%) | 27.67% (18.63%) |
| Unrelated | 78.33% (22.14%) | 84.00% (19.33%) |
| TOTAL | 56.25% (28.17%) | 57.75% (28.59%) |

These items were presented immediately following the EEG experimental task. Accuracy is separated based on medium of passage presentation.

A two-way repeated measures ANOVA was run to determine the statistical significance of the interaction between medium of presentation and accuracy across question types. No significant interaction was found (medium x item type: *F* (3, 147) = 0.89, *p* = .449), and the main effect of medium was also non-significant (*F* (1, 49) = 0.561, *p* = .457). However, the main effect of question type was significant (*F* (3, 147) = 85.105, *p* < .001), and planned comparisons (*t*-tests) revealed that accuracy for each of the question types was significantly different, in the following order from most to least accurate: Unrelated > Explicit (*t* (198) = 5.528, *p* < .001); > Paraphrase (*t* (192.19) = 3.586, *p* < .001); > Meaning Change (*t* (173.14) = 8.107, *p* < .001).

#### Delayed (Retention) Comprehension Task

In addition to collecting responses to the sentence verification items about each passage immediately following presentation, we asked participants to answer the same questions again within 24 hours after completing the lab session. However, the survey responses were accepted up to 168 hours (seven days) following the lab session. The goal was to gauge retention of the information presented in the passages, and to compare retention between media. Mean accuracy for each item type is presented below in Table 4.

**Table 4.** Mean percent correct responses for each sentence verification item type, delayed presentation.

| Sentence Verification | Digital Text Presentation | Print Text Presentation |
| --- | --- | --- |
| Item Type | % Correct (SD) | % Correct (SD) |
| Explicit | 63.33% (22.59%) | 63.33% (19.22%) |
| Paraphrase | 49.33% (20.75%) | 48.67% (23.77%) |
| Meaning Change | 24.67% (15.87%) | 25.33% (19.70%) |
| Unrelated | 63.33% (29.16%) | 65.67% (28.45%) |
| TOTAL | 50.17% (27.45%) | 50.75% (28.12%) |

These items were presented 1-7 days following the EEG experimental task. Accuracy is separated based on medium of passage presentation.

The pattern of responses to the delayed sentence verification task is similar to that of the immediate recall comprehension evaluation: meaning change items were responded to with the lowest accuracy, followed by paraphrase items. In this case, the accuracy for explicit and unrelated items appears equivalent, while overall accuracy is slightly lower for delayed vs. immediate evaluation. These results were confirmed with statistical analysis. A two-way repeated measures ANOVA was conducted, and no significant interaction between medium and item type was found (*F* (3, 147) = .195, *p* = .90). The main effect of medium was also non-significant (*p* = .75), but the main effect of item type was found to be significant (*F* (2.37, 115.91) = 41.240, *p* < .001). Planned comparisons (*t*-tests) showed that accuracy for unrelated and explicit items was not significantly different, but both were responded to significantly more accurately than paraphrase and meaning change items. Accuracy of responses to the paraphrase question type was greater than to the meaning change question type.

Additionally, we sought to identify significant differences between immediate recall comprehension (SVT items presented during the experiment run) and later retention accuracy (SVT items completed via online survey after at least 24 hours elapsed). Two separate two-way repeated measures ANOVAs were run, to observe the effects of time (immediate vs. delayed) and item type separately across the two mediums. For the digital passages, a significant interaction between time and item type was found (*F* (3, 132) = 3.204, *p* = .030), and the main effects of time (*F* (1, 44) = 13.02, *p* < .001) and item type (*F* (3, 132) = 49.697, *p* < .001) were also significant. Similarly, for the print passages, there was a significant interaction between time and item type (*F* (3, 132) = 5.448, *p* = .001) as well as significant main effects (time: *F* (1, 44) = 13.020, *p* < .001; item type: *F* (2.26, 99.64) = 60.197, *p* < .001). The effects of time reflected that total accuracy was significantly higher in the immediate responses to comprehension items than for the delayed responses (*t* (397.73) = 2.187, *p* = .03), while the interaction was driven by a difference in accuracy rates for the unrelated SVT items: on average 16.67% higher when responded to immediately after the passage reading task, compared to delayed responses (*t* (180.88) = 4.698, *p* < .001). The significant main effects of item type reflected that accuracy rates continued to follow the general pattern previously observed (Unrelated > Explicit: *t* (379) = 3.669, *p* < .001; > Paraphrase: *t* (392.61) = 5.814, *p* < .001; > Meaning Change: *t* (365.09) = 11.66, *p* < .001). With respect to delayed responses, the meaning change item type again yielded significantly fewer accurate responses than all other question types (Unrelated: *t* (165.42) = 11.699, *p* < .001; Explicit: *t* (192.3) = 13.85, *p* < .001; Paraphrase: *t* (189.09) = 8.433, *p* < .001). Paraphrase item types yielded significantly fewer accurate responses than the explicit and unrelated items (*t* (197.57) = -4.671, *p* < .001; *t* (186.27) = -4.273, *p* < .001, respectively). However, responses to the explicit and unrelated items did not differ with respect to accuracy (*t* (182.24) = 0.327, *p* = .744).

## Discussion

As alluded to above, this study took place against a complex background of research and environmental factors that contribute to the importance of the findings. The COVID pandemic was a time of unprecedented disruption to our educational systems, with as-yet little understood consequences for students. Amid pre-existing doubts about the impact of digital media on the development of reading and related skills, children were abruptly forced into online instruction and even more of their engagement with text, at all levels, now happens through various digital devices. These disruptions highlighted a challenge already being faced by educators: to understand how reading comprehension and learning are changing in the age of digital information. This investigation of the neural correlates of depth of processing during reading discourse across mediums in middle-school students is the first to apply event-related methodologies to this question, and is novel in its use of the N400 as an index. We drew upon the depth of processing theory introduced by Craik and Lockhart [28] to provide a theoretical framework for the investigation, alongside Kintsch’s [29, 30] view that text comprehension is a dynamic process of constructing meaning from semantic relations among words in the text and one’s stored knowledge about subject matter. We proposed that how readers engage with text/reading material may be a crucial determinant of differences in depth of processing for the semantic information contained in a text, consequently affecting the robustness of semantic memory structures that are established in support of reading comprehension. We extended the standard applications of the N400 to provide an index of processing depth associated with two mediums of text presentation: digital (via a laptop screen) and print (via a printed page).

We predicted that N400 responses to reading text presented in digital and print formats would differ. These predictions were largely supported by the data presented above. The waveforms indicate distinct brain responses across the two mediums. Consistent with our predictions, when passages were read on a laptop (digital), responses to subsequently presented words in the chimera (moderately related/moderately unrelated) category evoked activations similar to those associated with words that were unrelated to the text. This finding can be observed in the waveforms (Fig 3), and is supported by the lack of statistical significance in amplitude differences between chimera and unrelated word responses in the digital condition. The N400 waveforms in these two conditions can be observed to differ significantly from the response to related words.

In the print medium, we predicted that the N400 responses for the three conditions would be graduated with unrelated words producing the greatest negativity, the response to related words being the most attenuated, and responses to chimera words falling between. However, the N400 waveform patterned differently than expected (Fig 4). Mean amplitude values within the N400 time window were significantly different between related and unrelated words, and between chimera and unrelated words – consistent with our predictions. However, contrary to prediction, the amplitude differences in response to related and chimera stimuli were not significant.

Within the context of the depth of processing theory [28] the primary experimental manipulation in this study related to the chimera word stimuli. As prior behavioral studies have suggested, reading on a digital device promotes shallow reading. When classifying the chimera words, we stated that these stimuli could be perceived as related or unrelated given the center clustering of their word relatedness rating. Whether chimeras are perceived as related or unrelated to the text may depend on the strength of the encoded memory traces established during text discourse processing. Therefore, perception of chimera words as unrelated words would be consistent with shallow discourse processing as hypothesized in digital text reading, whereas chimera words perceived as related would be consistent with deeper discourse processing as observed in print reading.

Behaviorally, there was no distinction between classification of the chimera words by study participants following the digital or print presentations of texts; in both conditions, chimera words were most frequently identified as being “unrelated” to the text. The longer reaction times to related words than words in other probe conditions likely reflect response competition (e.g., [66]), and the patterning of reaction times between chimera and related words in the print condition, and between chimera and unrelated words in the digital condition, is an expected finding given the study prediction that more robust semantic networks were expected to develop following exposure to print vs. digital texts. However, observations of the ERP responses to chimera words provide a deeper insight.

The semantic judgement task prompted participants to decide whether presented probe words were related or unrelated, potentially shaping brain responses specific to the task at hand. Therefore, how deeply a participant read the text would likely contribute to whether they perceived the chimera word probes as either related or unrelated. This seems to bear out in the waveforms and statistics: responses to the chimera words track with responses to the unrelated words in the digital condition, and with responses to the related words in the print condition. The observed responses to the chimera word condition may index the robustness of context models for the text: if robust models are created, chimeras can be situated within the model affording greater processing efficiency, whereas when such words are situated within a less robust contextual model, as would be generated under shallower reading, the opposite would be expected. Under this interpretation, these ERP responses align with the study hypothesis and may indicate that a more “robust” semantic network was derived in response to texts presented in the print medium. Hence, we propose that the N400 brain responses observed are consistent with a finding of deeper text processing in print compared to digital media.

The increased use of digital materials alongside paper-based materials in learning environments has motivated many studies on the efficacy of reading and learning in one format versus the other (e.g., [1, 2, 5]). Investigations of reading comprehension and learning measured in terms of reading ability, reading rate, eye movement, and factual recall, have found no differences in student performance between working in the two mediums (e.g., [9, 14, 67, 68]). The present study is the first to evaluate depth of processing for print and digital informational texts in middle-school children using a brain measure (N400 ERP). Our findings contribute to this landscape by providing insights about the neurocognitive processing underlying reading comprehension. The study outcomes reveal differences in how the brain processes expository text when presented in digital and print mediums, with the former suggesting more shallow engagement and the latter conferring deeper engagement. This effect could indicate a “print advantage” with respect to depth of processing, in support of previous behavioral research [2].

### Study Limitations and Delimitations

As with any study that seeks to break new ground, there are important limitations to acknowledge and address in future work. Our study sample, despite our recruitment efforts, was skewed towards higher parental income and higher parental education levels and therefore does not adequately represent the diversity of the target populations (NYC metropolitan area). Future work should direct efforts towards recruitment of participants from a wider range of SES and parental educational backgrounds to determine whether the findings hold across demographic variables.

In addition, samples from communities without ready access to the internet would be important to evaluate since internet access and other amenities likely to predispose participants towards digital consumption of information may be lacking, so that students in such communities may be less experienced or less prepared to read texts digitally. This could lead to different patterns of reading preference, experience, and relative advantage; for example, less familiarity with digital media could be associated with less robust semantic memory structures established for information presented in this medium, therefore resulting in lower processing efficiency.

Our participants were middle-school children in the New York City metropolitan area, mostly reporting post-secondary parental education and mid-to-high SES backgrounds. Our entire sample was born after 2010, and so all can be considered “digital natives” or members of “iGen” (in the sense defined by Twenge et al. [69]). This strongly suggests that digital exposure would have been optimal for these participants throughout their lives, predisposing them to be expert consumers of text and other kinds of information in digital formats. It is also possible that our sample may have been taught or absorbed strategies for reading and learning online given the prevalence of online schooling in New York City during the pandemic that preceded our data collection. Within the current sample, there were no significant differences between the medium of presentation in comprehension of the texts, reading times, or performance on a measure of information retention. Nonetheless, the N400 effects remain; while our findings suggest differences in the efficiency of neurocognitive processing across different media, further research is needed. Overall, the underlying nature of the interaction between experience with particular media and reading comprehension remains to be addressed.

Despite earlier debates about the context of digital adaptations in learning and differences in access to digital media (summarized by Evans & Robertson [70]), iGen access and exposure to digital media appears uniform across gender, race/ethnicity, and socioeconomic status [69] – even leading to concerns that there has been a displacement of so-called “legacy media” (a term encompassing everything from print books and magazines to television). Carr [71] and Wolf [72] have also suggested that the seemingly shallow processing associated with accessing texts in digital formats could relate to readers being primed by the larger culture of the digital age, to access information in smaller “bits” and to process it less deeply when reading from a screen. Despite such concerns, the majority of our sample identified a preference for print over digital media (similar to that observed by Kretzschmar et al. [14]), and we observed a corresponding print advantage in the N400 data for semantic processing of text-related concepts.

Our study parameters were necessarily delimited in many ways. We selected middle-school children for our cross-sectional study design, to reflect the age at which brain adaptations for successful attainment of reading skills are considered to be underway [73, 74]. Chall [75] identified our selected age range as critical in reading development, having proposed a shift in fourth grade from “learning to read” to “reading to learn” – based on the proposal that early learning of basic reading-related skills (such as grapheme-to-phoneme correspondences) shifts around this age to higher-level skills including reading comprehension. Hence, considerations of earlier stages in reading development, and how these adaptations interact with exposure to texts in different mediums, limit the generalizability of our findings.

Other neurophysiological approaches to understanding reading development provide evidence to suggest that a focus on older age groups could also be relevant for future work. For example, Coch [76] used the N400 to investigate orthographic, semantic, and phonological processing in children from 3rd-5th grade, as well as college-age students. Participants were presented with real words, pseudowords, non-pronounceable letter strings, false font strings, and animal names. While an adult-like response was observed for stimuli tapping into semantic and phonological processing, the child participants (but not the college students) showed responses to false font strings similar to their word reading responses. Coch proposed that this changes by adulthood due to extensive reading experience and fine-tuned word processing; but it is not clear at what age automaticity might be attained and what specific neural processes might index such attainment. Until recently, there has been a paucity of evidence-based support for pedagogical practice and policy (e.g., [77]); hence, there is a need to evaluate the application of neurophysiological measures to support effective approaches to developing skilled deep readers.

Another study limitation is instantiated in the limited number of standardized measures conducted to ensure that participants were typically developing readers for their grade and age. Time constraints related to the anticipated average attention span of our target population prohibited the inclusion of other potentially valuable measures. In the future, measures of vocabulary and reading experience could offer deeper insights regarding individual differences. Additionally, we generated recall and retention comprehension question as one measure to ensure equivalency across passages. Unfortunately, missing data from both the recall and retention assessments, compounded by the fact that there were only two items for each question type, made comparisons with the N400 mean amplitude measure difficult.

During our development of the text passages used as stimuli in this study, we made a decision to work with expository or informational texts. This decision was based on meta-analyses [1, 2] showing that reading performance advantages when reading printed text on paper versus digital formats held for expository and informational texts but not narrative texts. The selection of expository text allowed us to more effectively control propositional counts for each passage, and to develop passages similar to those likely encountered by children in their learning environments. However, it is possible that distinct effects on indices of neural engagement, and/or behavioral indices of comprehension, could be identified if the texts were narrative in nature. Comparisons between responses to matched sets of narrative and expository texts would be valuable in future work.

## Conclusions

As we have described here, this study marks the first step towards systematic application of neurophysiological methods to understand the implications and neural underpinnings of reading in print vs. digital media, at a crucial stage in literacy acquisition. An important question raised by these findings concerns the implications for classroom instruction of reading and learning via paper-based texts compared to texts delivered on digital platforms. The question is particularly relevant given the near ubiquitous use of digital platforms for delivery of instruction and information at school and at home.

For reasons related to study delimitations and limitations we think it too early to generate a set of recommendations for adaptation in the classroom. However, we do think that these study outcomes warrant adding our voices to those of Delgado et al. [2] in suggesting that we should not yet throw away printed books, since we were able to observe in our participant sample an advantage for depth of processing when reading from print. Applications for digital reading should not be dismissed, either: the observation of a potential print advantage does not negate the value of rapid access to information that could be supported by digital reading. It may be that classroom practices should strategically match reading strategies and mediums to task, such that printed media are employed when deeper processing is required while digital access to text is utilized for other needs.

Another reason not to dismiss digital reading platforms is their potential to benefit children with reading disabilities. Research in this area suggests that digital reading strategies may be utilized in support of reading proficiency [78] and comprehension [79] in this population. However, reading disabilities are vastly heterogeneous, and there are concomitant difficulties with identification (e.g., [80]), alongside a corresponding array of interacting causal mechanisms that need to be described at multiple levels – at least, behaviorally, neurophysiologically, and genetically (e.g., [81]). Hence, further investigations of the effectiveness of digital and print text presentations for dyslexia and other reading disabilities will be needed.

## Acknowledgements

A special thanks to the incredible research team from the Neurocognition of Language Lab at Teachers College, including Sarah Bennett, M.S., Girija Chatufale, M.S., Kenny Fabara, M.S., Kai Gilchrist, M.S., Zeina Haidar, M.S., Luiza Lodder, M.S., Emily Mantaro, M.S., Michela Thomsen, M.S., and Huiyu (Bonnie) Yang, M.S. We also thank Dakota Egglefield, Ph.D., for her contributions to every aspect of our behavioral data collection; and Alexa Monachino for support during data analysis applications. We want to thank Dr. Kara Federmeier for her invaluable suggestions on earlier versions of this article, and Dr. Trey Avery for his continued support with the technology and the data. This research would not be possible without the wonderful families and participants who let us borrow their magnificent brains – our deep appreciation for their interest, support, and time.

## References

1. Clinton V. Reading from paper compared to screens: A systematic review and meta-analysis. J Res Read. 2019 Jan 13;42(2):288–325. 10.1111/1467-9817.12269

2. Delgado P, Vargas C, Ackerman R, Salmerón L. Don’t throw away your printed books: A meta-analysis on the effects of reading media on reading comprehension. Educ Res Rev. 2018 Nov;25:23–38. 10.1016/j.edurev.2018.09.003

3. Rideout VJ, Foehr UG, Roberts DF. Generation M2: Media in the lives of 8-to 18-year-olds. Henry J. Kaiser Family Foundation; 2010.

4. Sidi Y, Shpigelman M, Zalmanov H, Ackerman R. Understanding metacognitive inferiority on screen by exposing cues for depth of processing. Learn Instr. 2017 Oct;51:61–73. 10.1016/j.learninstruc.2017.01.002

5. Singer LM, Alexander PA. Reading across mediums: Effects of reading digital and print texts on comprehension and calibration. J Exp Educ. 2016 Mar 9;85(1):155–72. 10.1080/00220973.2016.1143794

6. DeJong MT, Bus AG. The efficacy of electronic books in fostering kindergarten children’s emergent story understanding. Read Res Q. 2004 Oct 12;39(4):378–93. 10.1598/rrq.39.4.2

7. Dündar H, Akçayır M. Tablet vs. paper: The effect on learners’ reading performance. International Electronic J Elem Educ. 2012 Jan 1;4(3):441–50.

8. Kong Y, Seo YS, Zhai L. Comparison of reading performance on screen and on paper: A meta-analysis. Comput Educ. 2018 Aug;123:138–49. 10.1016/j.compedu.2018.05.005.

9. Margolin SJ, Driscoll C, Toland MJ, Kegler JL. E-readers, computer screens, or paper: Does reading comprehension change across media platforms? Appl Cogn Psychol. 2013 May 28;27(4):512–9. 10.1002/acp.2930.

10. Nichols M. Reading and studying on the screen: An overview of literature towards good learning design practice. J Open Flex Dist Learn. 2016 Aug 25;20(1):33–43. 10.3316/informit.195571684952519

11. Noyes JM, Garland KJ. Computer-vs. paper-based tasks: Are they equivalent? Ergonom. 2009;51:1352–75. 10.1080/00140130802170387.

12. Singer LM, Alexander PA. Reading on paper and digitally: What the past decades of empirical research reveal. Rev Educ Res. 2017 Jul 21;87(6):1007–41. 10.3102/0034654317722961

13. Wang S, Jiao H, Young MJ, Brooks T, Olson J. Comparability of computer-based and paper-and-pencil testing in K–12 reading assessments. Educ Psychol Meas. 2007 Sep 12;68(1):5–24. 10.1177/0013164407305592.

14. Kretzschmar F, Pleimling D, Hosemann J, Füssel S, Bornkessel-Schlesewsky I, Schlesewsky M. Subjective impressions do not mirror online reading effort: Concurrent eeg-eyetracking evidence from the reading of books and digital media. PLoS ONE. 2013 Feb 6;8(2):e56178. 10.1371/journal.pone.0056178

15. Lenhard W, Schroeders U, Lenhard A. Equivalence of screen versus print reading comprehension depends on task complexity and proficiency. Discourse Process. 2017 May 4; 54(5-6):427–45. 10.1080/0163853x.2017.1319653

16. Singer LM, Alexander PA, Berkowitz LE. Effects of processing time on comprehension and calibration in print and digital mediums. Int J Exp Educ. 2017 Dec 21;87(1):101–15. 10.1080/00220973.2017.1411877.

17. Connell C, Bayliss L, Farmer W. Effects of e-book readers and tablet computers on reading comprehension. Int J Instruc Media. 2012;39(3):131–40.

18. Daniel DB, Woody WD. E-textbooks at what cost? Performance and use of electronic v. print texts. Comput Educ. 2013 Mar;62:18–23. 10.1016/j.compedu.2012.10.016.

19. Kerr MA, Symons SE. Computerized presentation of text: effects on children’s reading of informational material. Read Writ. 2006 Feb;19(1):1–19. 10.1007/s11145-003-8128-y

20. Kim HJ, Kim J. Reading from an LCD monitor versus paper: Teenagers’ reading performance. Int J Res Stud Educ Technol. 2013 Apr 5;2(1). 10.5861/ijrset.2012.170

21. Deschryver M, Spiro RJ. New forms of deep learning on the web: Meeting the challenge of cognitive load in conditions of unfettered exploration in online multimedia environments. In: Zheng R, editor. Cognitive effects of multimedia learning. IGI Global eBooks; 2009. p. 134–52. 10.4018/978-1-60566-158-2.ch008

22. Ackerman R, Lauterman T. Taking reading comprehension exams on screen or on paper? A metacognitive analysis of learning texts under time pressure. Comput Hum Behav. 2012 Sep;28(5):1816–28. 10.1016/j.chb.2012.04.023.

23. Wästlund E, Reinikka H, Norlander T, Archer T. Effects of VDT and paper presentation on consumption and production of information: Psychological and physiological factors. Comput Hum Behav. 2005 Mar;21(2):377–94.

24. Lauterman T, Ackerman R. Overcoming screen inferiority in learning and calibration. Comput Hum Behav. 2014 Jun;35:455–63. 10.1016/j.chb.2014.02.046

25. Singer Trakhman LM, Alexander PA, Berkowitz LE. Effects of processing time on comprehension and calibration in print and digital mediums. J Exp Educ. 2017 Dec 21;87(1):101–15. 10.1080/00220973.2017.1411877

26. Mangen A, Walgermo BR, Brønnick K. Reading linear texts on paper versus computer screen: Effects on reading comprehension. Int J Educ Res. 2013 Jan;58(58):61–8. 10.1016/j.ijer.2012.12.002

27. Eshet-Alkalai Y, Geri N. Does the medium affect the message? The influence of text representation format on critical thinking. Hum Sys Manag. 2007 Dec 20;26(4):269–79. 10.3233/hsm-2007-26404

28. Craik FIM, Lockhart RS. Levels of processing: A framework for memory research. J Verbal Learning Verbal Behav. 1972 Dec;11(6):671–84.

29. Kintsch W. Comprehension: a paradigm for cognition. Cambridge: Cambridge Univ. Press; 1998.

30. Kintsch W. Psychological models of reading comprehension and their implication for assessment. In: Sabatino J, Albro E, O’Reilly T, editors. Measuring up: Advances in how we assess reading ability. Lanham, MD: Rowman and Littlefield Publishers, Inc.; 2012. p. 21–38.

31. Craik FIM, Tulving E. Depth of processing and the retention of words in episodic memory. J Exp Psychol: Gen. 1975;104(3):268–94. 10.1037/0096-3445.104.3.268

32. Tulving E. Episodic and semantic memory. In: Tulving E, Donaldson W, editors. Organization of memory. Cambridge, MA: Academic Press; 1972. p. 381–403.

33. Anderson J, Reder L. An elaborative processing explanation of depth of processing. In: Cermak L, Craik F, editors. Levels of processing in human memory. Mahwah, NJ: Lawrence Earlbaum Associates; 1979. p. 385–404.

34. Federmeier KD, Laszlo S. Time for meaning: Electrophysiology provides insights into the dynamics of representation and processing in semantic memory. In: Ross B, editor. The psychology of learning and motivation. Cambridge, MA: Elsevier Academic Press; 2009. p. 1–44. 10.1016/S0079-7421(09)51001-8

35. Kutas M, Federmeier KD. Thirty years and counting: Finding meaning in the N400 component of the event-related brain potential (ERP). Ann Rev Psychol. 2011 Jan 10;62(1):621–47. 10.1146/annurev.psych.093008.131123

36. Kincaid J, Fishburne R, Rogers R, Chissom B. Derivation of new readability formulas (Automated Readability Index, Fog Count, and Flesch Reading Ease formula) for Navy-enlisted personnel. Research Branch Report 8-75: Chief of Naval Technical Training; 1975.

37. Gunning R. The technique of clear writing. New York: Mcgraw-Hill Book Co; 1952.

38. McLaughlin G. SMOG grading: A new readability formula. J Read. 1969;12(8):639–46.

39. Bovair S, Kieras D. A guide to propositional analysis for research on technical prose. Technical Report No. 8, University of Arizona; 1981.

40. Kintsch W. The representation of meaning in memory. Hillsdale, NJ: Lawrence Erlbaum Associates; 1974.

41. Turner A, Greene E. The construction and use of a propositional text base. Institute for the Study of Intellectual Behavior Technical Report No. 63, University of Colorado; 1977.

42. Royer JM. Developing reading and listening comprehension tests based on the Sentence Verification Technique (SVT). J Adol Adult Literacy. 2001 Sep 1;45(1):30–41. https://www.jstor.org/stable/40007629

43. Kuperman V, Stadthagen-Gonzalez H, Brysbaert M. Age-of-acquisition ratings for 30,000 English words. Beh Res Meth. 2012 Dec;44:978–90.

44. Fellbaum C. A semantic network of English: the mother of all WordNets. Comp Humanities. 1998 Mar;32:209–20.10.1023/A:1001181927857

45. Miller GA. WordNet: a lexical database for English. Commun ACM. 1995 Nov 1;38(11):39–41.

46. Princeton University. WordNet (Version 3.0). 2010. https://wordnet.princeton.edu/

47. Brysbaert M, New B. Moving beyond Kučera and Francis: A critical evaluation of current word frequency norms and the introduction of a new and improved word frequency measure for American English. Beh Res Meth. 2009 Nov;41(4):977–90. 10.3758/brm.41.4.977

48. Coltheart M. The MRC psycholinguistic database. Q J Exp Psychol. 1981 Nov;33(4):497–505. 10.1080/14640748108400805

49. Brysbaert M, Warriner AB, Kuperman V. Concreteness ratings for 40 thousand generally known English word lemmas. Beh Res Meth. 2014 Sep;46:904–11. 10.3758/s13428-013-0403-5

50. Dubossarsky H, De Deyne S, Hills TT. Quantifying the structure of free association networks across the life span. Dev Psychol. 2017 Aug;53(8):1560. 10.1037/dev0000347

51. Lindsay BG. Mixture models: Theory, geometry, and applications. NSF-CBMS; 1995.

52. McLachlan G, Peel, D. Finite mixture models. Hoboken, NJ: John Wiley & Sons; 2000.

53. Wechsler D. Wechsler intelligence scale for children (5th ed.). New York, NY: Pearson; 2014.

54. Woodcock, R.W. Woodcock reading mastery tests (3rd ed.). New York, NY: Pearson; 2011.

55. Swanson HL. Generality and modifiability of working memory among skilled and less skilled readers. J Educ Psychol. 1992;84(4):473–88. 10.1037/0022-0663.84.4.473

56. Gabard-Durnam LJ, Mendez Leal AS, Wilkinson CL, Levin AR. The Harvard Automated Processing Pipeline for Electroencephalography (HAPPE): Standardized processing software for developmental and high-artifact data. Front Neurosci. 2018 Feb 27;12. 10.3389/fnins.2018.00097

57. Monachino AD, Lopez KL, Pierce LJ, Gabard-Durnam LJ. The HAPPE plus Event-Related (HAPPE+ ER) software: A standardized preprocessing pipeline for event-related potential analyses. Dev Cogn Neurosci. 2022 Oct 1;57:101140. 10.1101/2021.07.02.450946

58. Šoškić A, Jovanović V, Styles SJ, Kappenman ES, Ković V. How to do better N400 studies: reproducibility, consistency and adherence to research standards in the existing literature. Neuropsychol Rev. 2022 Sep;32(3):577–600. 10.31234/osf.io/jp6wy

59. Chwilla DJ, Brown CM, Hagoort P. The N400 as a function of the level of processing. Psychophysiol. 1995 May;32(3):274–85. 10.1111/j.1469-8986.1995.tb02956.x

60. Hald LA, Steenbeek-Planting EG, Hagoort P. The interaction of discourse context and world knowledge in online sentence comprehension. Evidence from the N400. Brain Res. 2007 May 18;1146:210–8. 10.1016/j.brainres.2007.02.054

61. Kutas M. In the company of other words: Electrophysiological evidence for single-word and sentence context effects. Lang Cogn Proc. 1993 Nov 1;8(4):533–72. 10.1080/01690969308407587

62. Nieuwland MS, Van Berkum JJ. When peanuts fall in love: N400 evidence for the power of discourse. J Cogn Neurosci. 2006 Jul 1;18(7):1098–1111. 10.1162/jocn.2006.18.7.1098

63. Nieuwland MS, Otten M, Van Berkum JJ. Who are you talking about? Tracking discourse-level referential processing with event-related brain potentials. J Cogn Neurosci. 2007 Feb 1;19(2):228–36. 10.1162/jocn.2007.19.2.228

64. George MS, Mannes S. Global semantic expectancy and language comprehension. J Cogn Neurosci. 1994 Jan 1;6(1):70–83. 10.1162/jocn.1994.6.1.70

65. Van Berkum JJ, Brown CM, Zwitserlood P, Kooijman V, Hagoort P. Anticipating upcoming words in discourse: evidence from ERPs and reading times. J Exp Psychol: Learn Mem Cogn. 2005 May;31(3):443. 10.1037/0278-7393.31.3.443

66. Lupker SJ. The semantic nature of response competition in the picture-word interference task. Mem Cogn. 1979 Nov;7(6):485–95.

67. Bodmann SM, Robinson DH. Speed and performance differences among computer-based and paper-pencil tests. J Educ Comp Res. 2004 Jul;31(1):51–60. 10.2190%2Fgrqq-yt0f-7lkb-f033

68. Van de Velde C, von Grunau M. Tracking eye movements while reading: Printing press versus the cathode ray tube. Percept 2003 Jan;32:107.

69. Twenge JM, Martin GN, Spitzberg BH. Trends in US Adolescents’ media use, 1976– 2016: The rise of digital media, the decline of TV, and the (near) demise of print. Psychol Pop Media Cult. 2019 Oct;8(4):329. 10.1037/ppm0000203

70. Evans C, Robertson W. The four phases of the digital natives debate. Hum Beh Emerg Technol. 2020 Jul;2(3):269–77. 10.1002/hbe2.196

71. Carr N. The shallows: What the Internet is doing to our brains. New York, NY: WW Norton & Company; 2020 Mar 3.

72. Wolf M. Reader come home: The reading brain in a digital world. New York, NY: Harper Collins; 2018.

73. Dehaene S, Cohen L. The unique role of the visual word form area in reading. Trends Cogn Sci. 2011 Jun 1;15(6):254–62. 10.1016/j.tics.2011.04.003

74. McCandliss BD, Cohen L, Dehaene S. The visual word form area: expertise for reading in the fusiform gyrus. Trends Cogn Sci. 2003 Jul 1;7(7):293–9. 10.1016/s1364-6613(03)00134-7

75. Chall J. S. Stages of reading development. New York, NY: McGraw-Hill; 1983.

76. Coch D. The N400 and the fourth grade shift. Dev Sci. 2015 Mar;18(2):254–69. 10.1111/desc.12212

77. Slavin RE. How evidence-based reform will transform research and practice in education. Educ Psychol. 2020 Jan 2;55(1):21–31. 10.1080/00461520.2019.1611432

78. Gunter GA, Kenny RF. UB the director: Utilizing digital book trailers to engage gifted and twice-exceptional students in reading. Gift Educ Int. 2012 May;28(2):146–60. 10.1177/0261429412440378

79. Ertem IS. The effect of electronic storybooks on struggling fourth-graders’ reading comprehension. Turkish Online J Educ Technol. 2010 Oct;9(4):140–55.80.

80. Wagner RK, Zirps FA, Edwards AA, Wood SG, Joyner RE, Becker BJ, Liu G, Beal B. The prevalence of dyslexia: A new approach to its estimation. J Learn Dis. 2020 Sep;53(5):354–65. 10.1177/0022219420920377

81. Centanni T. Neural and genetic mechanisms of dyslexia. In: Argyropoulos G, editor. Translational neuroscience of speech and language disorders. New York, NY: Springer; 2020. p. 47–65. 10.1007/978-3-030-35687-3_4

